# Physiological load via Voluntary Wheel Running maintains Achilles tendon homeostasis by region-specific cellular responses

**DOI:** 10.64898/2026.05.28.722285

**Authors:** Samantha N. Muscat, Elsa Lecaj, Nolan Sparks, Alexander Kollar, Eowyn Kroening, Mark Buckley, Anne E.C. Nichols

**Author notes:** **Corresponding Author:** Anne E.C. Nichols, PhD, 601 Elmwood Avenue, Box 665G Rochester, NY 14642.

## Abstract

Physiological load is vital for maintaining tendon homeostasis, preserving the organized extracellular matrix that enables tendons to withstand extreme forces. Although tenocytes are regarded as the primary regulators of extracellular matrix production, precisely how cells facilitate the maintenance of homeostasis in response to physiological load is poorly understood. Here, we used Voluntary Wheel Running (VWR) as a model of physiological load to delineate the specific cellular contributions to mouse Achilles tendon homeostasis. Eight weeks of VWR led to a smaller cross-sectional area, increased mechanical and material properties at the midsubstance, which corresponded to a decreased proportion of small (0-60 nm) collagen fibrils and an increased proportion of larger (100-60 nm) collagen fibrils compared to sedentary controls. Using Visium HD spatial transcriptomics, we identified region-specific cell clusters (insertion vs. midsubstance). In response to physiological load, cells in the insertion and midsubstance upregulate distinct genes that reinforce the fibrocartilage interface and collagen-rich tendon matrix, respectively. Notably, *Clu*, *Myoc,* and *Ccdc80* were upregulated with VWR in the midsubstance, with previously uncharacterized roles in tendon homeostasis. Together, our findings suggest that in response to physiological load, tendon cells maintain homeostasis by region-specific responses. Given that insertional and midsubstance tendinopathy is function-limiting and painful, defining the region-specific cellular responses will be key to advancing therapeutic prospects for tendon health.

**New and Noteworthy:** This study is the first spatially rigorous characterization of the tendon response to physiological load using a Voluntary Wheel Running (VWR) model. VWR led to smaller, stronger, but not stiffer tendons at the midsubstance compared to sedentary controls. This corresponded with significant decreased proportion of small collagen fibrils and a shift toward an increased proportion of large collagen fibrils. Using Visium HD spatial transcriptomics, we identified region-specific transcriptional responses to physiological load that maintain homeostasis.

**Graphical Abstract:** 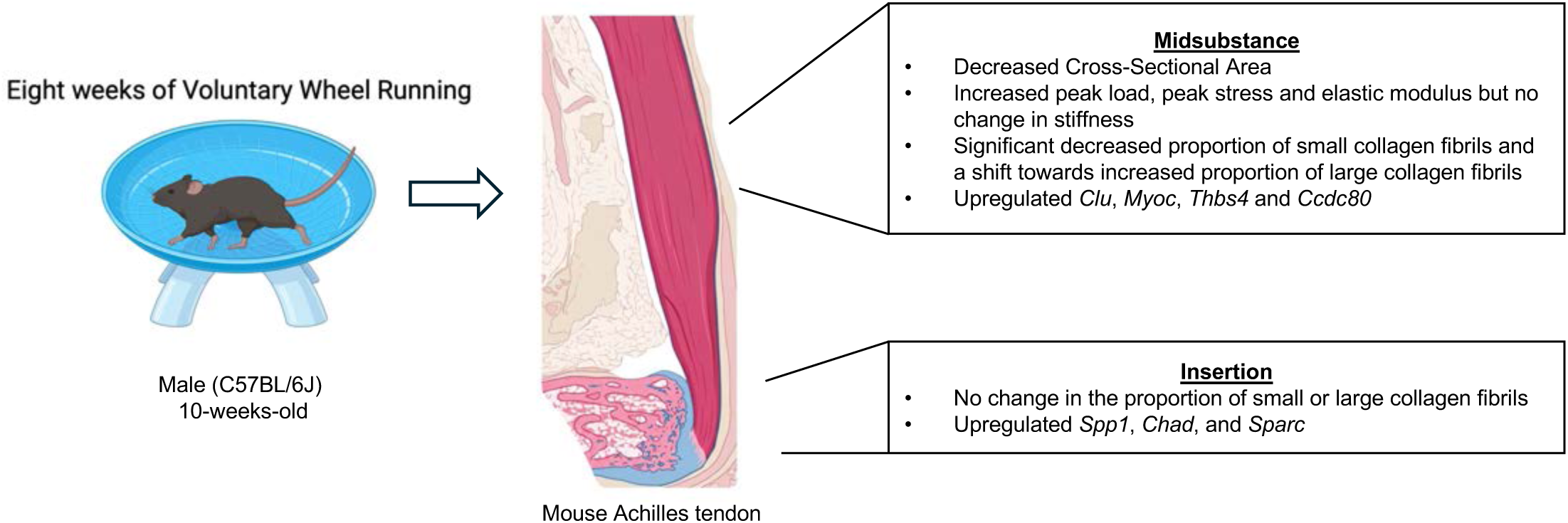

## Introduction

Tendons are soft tissue bridges between muscle and bone that transmit forces, facilitating successful musculoskeletal movement. The tendon is primarily composed of type 1 collagen fibers, which self-assemble into a hierarchical, parallel arrangement that supports high tensile strength (1). Non-collagenous extracellular matrix (ECM) proteins such as proteoglycans (1, 2) play a direct role in collagen fiber formation, and their unique ability to bind water molecules provides shock absorption during movement (2). The tendon’s highly organized ECM structure enables it to withstand high loads (3). For example, during moderate exercise (walking or running), the human Achilles tendon experiences load up to eight times one’s body weight (4). This physiological load is required to maintain the tendon’s organized structure and biomechanical properties, otherwise known as tendon homeostasis (5–7). Indeed, both over- and under-loading can lead to tendon degeneration and loss of function (8). Tenocytes, specialized fibroblasts within the tendon body, are the primary cellular mediators of mechanical loading (9, 10). In response to mechanical loading *in vitro,* tenocytes upregulate important ECM-related genes, including type 1 collagen (*Col1a1*, *Col1a2*) (11–14), matrix metalloproteinases and their inhibitors (*Mmp1*, *Timp1*) (12, 15, 16), and proteoglycans (*Dcn* and *Fbn*) (11), which are thought to promote tendon homeostasis by repairing, maintaining, and reinforcing the matrix. *In vivo* studies in both humans and animals have demonstrated that exercised tendons are stronger (increased biomechanical properties, including stiffness and peak load) than unexercised tendons (17, 18), and that increased strength is associated with increased collagen amount (% collagen dry weight) (19, 20) and size (cross-sectional area [CSA]) (21, 22). However, our current understanding of the tendon response to physiological load is limited by considerable variability, with the results of *in vivo* studies depending on tendon type, age, loading model, and time of evaluation. Moreover, the cellular mechanisms underlying these responses have been characterized largely using *in vitro/ex vivo* systems that do not fully recapitulate the *in vivo* mechanical environment.

Crucially, our understanding of cellular homeostatic maintenance is hindered by the paucity of animal models of physiological loading, as most focus on promoting tendon degeneration to model tendinopathy (8). For example, forced treadmill running (FTR) is commonly used to induce tendinopathy in Achilles (11, 23–27) and supraspinatus (28–30) tendons, as evidenced by disorganized collagen matrix and decreased mechanical properties, by forcing mice/rats to run under adverse conditions (electrical shock, blasts of air, or manual prodding). This induces a significant stress response as indicated by reports of increased anxiety and corticosterone (31, 32), which can result in confounding effects (33). Voluntary Wheel Running (VWR), however, does not require a negative stimulus, as it takes advantage of a mouse’s natural inclination to run (34). VWR also recapitulates normal running behavior in mice and does not interfere with their nocturnal activity (35). Indeed, VWR promotes adaptations consistent with moderate exercise in cardiac and skeletal muscle (36). Importantly, VWR has been shown to attenuate disease symptoms in a mouse model of colitis, while FTR exacerbated disease symptoms and increased mortality rates (37). Taken together, these studies suggest that the confounding effects from FTR induce systemic stress, which masks true exercise behaviors, while VWR mimics beneficial moderate exercise.

Thus, in the present study, we used VWR to understand how mouse Achilles tendons maintain homeostasis in response to physiological load. We demonstrate that eight weeks of VWR results in decreased CSA and increased mechanical and material properties at the midsubstance, corresponding with a significant decrease in the proportion of small collagen fibrils and a shift towards a larger proportion of larger collagen fibrils. Given the changes in biomechanical properties and collagen fibril size at the midsubstance, we leveraged spatial transcriptomics to define the precise cellular processes that maintain homeostasis in response to physiological load in a spatially-dependent manner. Together, these findings provide the first comprehensive characterization of how physiological loading affects tendons and reveal region-specific cellular responses.

## Methods

### Mice

This study was carried out in strict accordance with the recommendations in the *Guide for the Care and Use of Laboratory Animals* of the National Institutes of Health (Bethesda, MD, USA). All animal procedures and experiments were approved by the University Committee on Animal Research (UCAR) at the University of Rochester. Ten-week-old male C57BL/6J mice (#000664) were purchased from Jackson Laboratory. For all experiments, mice were singly housed to ensure unimpeded access to running wheels and to facilitate continuous recording of running behavior for individual mice. For this study, male mice were used in all experiments because they do not exhibit a significant stress response when singly housed (38). At the end of the running experiment, mice were euthanized by carbon dioxide inhalation followed by cervical dislocation unless otherwise noted.

### Voluntary Wheel Running (VWR) Setup

VWR mice (n=20) were allowed to run for eight weeks on the open surface of a slanted plastic saucer-shaped wheel placed inside the mouse cage (Med Associates, Inc. #ENV-047). Sedentary controls (n=20) had wheels locked to prevent running. Wheel revolutions were wirelessly transmitted to a hub so that the frequency and rate of running could be captured continuously throughout the experiment. Distance (km) was calculated by the total number of wheel revolutions and the radius of the wheel (Wheel Manager Data Acquisition Software, Med Associates, Inc.). Data were binned by hour to calculate the weekly and total running distance for individual mice. One VWR mouse was excluded from analysis due to natural death after 6 weeks of running.

### Biomechanical Assessment

Following euthanasia, the hindlimbs were cut at the patellar tendon, and the skin was carefully dissected. Hindlimbs were stored in a petri dish covered with gauze and PBS at -20°C until testing. Hindlimbs were thawed overnight at 4°C, and then the Achilles tendons were carefully dissected from hindlimbs in both groups (n=10 per group). Briefly, the plantaris tendon was removed, and the gastrocnemius muscle was carefully dissected from its proximal end. The hind paw was cut off with the calcaneal insertion still attached to the Achilles tendon. Custom-designed grips were 3D printed (Original Prusa MK4, Prusa Research) using polylactic acid. Sandpaper was glued onto the grips to minimize slipping of the tendon. Two grips were placed in a custom 3D printed alignment tool to ensure a consistent gauge length of 4.35 mm. Achilles tendons were placed onto the sandpaper on the grips with superglue. An additional two grips are then carefully laid on the tendon and secured with button head screws. Gripped tendons were imaged under a custom plexiglass platform to measure tendon width and thickness. Three measurements of tendon width and thickness were used to calculate the average CSA for each sample. Gripped tendons underwent tensile testing using a customized uniaxial micro tester (eXpert 4000 Microtester and MTESTQuattro software). Tendons were preloaded to 0.1N to remove slack, preconditioned with 10 cycles at 6% strain, 500-second stress relaxation at 8% and 16% strain, and then underwent a single ramp-to-failure (1 mm/s) at the midsubstance. Force-displacement curves were plotted and analyzed to determine stiffness and peak load. Stiffness was determined by the slope of the linear portion of the force-displacement curve.

Peak load was determined by the maximum load at failure. Stiffness and peak load were divided by the CSA to determine the elastic modulus and peak stress, respectively. Stress relaxation ratio (S: R) was determined as the relative decrease in tissue stress from the start of the step until the end of the relaxation period and normalized to its starting value. Time to 50% relaxation was determined as the number of seconds required for tissue stress to decrease to 50% of the total stress relaxation response.

### Transmission Electron Microscopy (TEM) Imaging and Quantification

Hindlimbs were dissected at the patellar tendon and immediately immersion-fixed in 3% glutaraldehyde/3% formaldehyde in 0.1M sodium cacodylate buffer (Electron Microscopy Sciences #15950) with 0.05% tannic acid for 24 hours at 4°C (n=5). Following fixation, the gastrocnemius muscle with the Achilles tendon attached was carefully teased to separate it from the hindlimb. The plantaris tendon was removed. The hind paw was then separated by carefully cutting around the calcaneus to ensure that the Achilles tendon remained attached. Surrounding fat from the Kager’s fat pad was carefully removed. For the insertion, the Achilles tendons were cut 2 mm from the point at which the tendons insert into the calcaneus. An additional 1 mm of tissue was removed to avoid overlap with the midsubstance, and the remaining tendon was used to analyze the midsubstance. Samples were then sent to the Shriners Children’s Hospital Micro-Imaging Core (Portland, OR) for processing and imaging. Briefly, samples were washed with DMEM, then decalcified in 0.2M EDTA with 50 mM TRIS pH 7.8 in a laboratory microwave (Ted Pella Inc.), operated at 97.5 watts for twenty 99-minute cycles. Samples were then fixed again in 3% glutaraldehyde/3% formaldehyde with 0.05% tannic acid overnight. After overnight fixation, samples were rinsed in DMEM and then post-fixed in 1% osmium tetroxide at 4°C. Samples were dehydrated and infiltrated to 100% Spurr’s epoxy with the aid of microwave energy, followed by an additional overnight in Spurr’s epoxy at 4°C before polymerization at 70°C. Transverse sections (60 nm) were contrasted with uranyl acetate and lead citrate and imaged on an FEI G20 transmission electron microscope operated at 120 kV with an AMT 2K x 2K camera. Insertion and midsubstance portions were divided into a 9-square grid system to ensure uniform coverage of the entire tendon. Three images were taken from each region at 50,000X magnification. A straight line was drawn along the x-axis of each fibril within each image in ImageJ software to determine fibril diameter. Only fibrils exhibiting a complete circumference were included in the measurements. Fibril density was calculated as fibrils per µm^2^.

### Collagen Hybridizing Peptide Staining

Hindlimbs were cut at the patellar tendon to ensure proper tension from the gastrocnemius muscle during fixation (n=5). Skin and toes were carefully removed to allow successful penetration of the fixative and decalcifying solutions. Hindlimbs were immediately immersion-fixed in 10% neutral buffered formalin for 24 hours, decalcified in Webb-Jee 14% EDTA solution (39) for 4 days, and immersed in 30% sucrose overnight for cryoprotection, all at 4°C. Hindlimbs were cut as described above and embedded in Frozen Section Media (Leica #3801480). Sagittal sections (10 uM) were cut using a cryostat using a cryotape-transfer method (40). Frozen sections were mounted on glass slides using 1% chitosan in 0.25% acetic acid and allowed to dry overnight at 4°C. Collagen hybridizing peptide-Cy3 (CHP) solution (3Helix #RED60) was diluted to 20 µM in PBS. The diluted CHP solution was incubated in a water bath at 80°C to dissociate triple helices into monomers. Following water bath incubation, 50 µL of CHP solution was added to frozen sections and allowed to incubate overnight at 4°C. The following day, frozen sections were washed with PBS for 5 minutes. Nuclei were counterstained with Hoechst 33342 (NucBlue™ Live ReadyProbes™ Reagent, Thermo Fisher Scientific), and slides were cover slipped (ProLong™ Diamond Antifade Mountant, Thermo Fisher Scientific). Imaging was performed using a VS120 Virtual Slide Microscope (Olympus).

### Paraffin Histology

Hindlimbs were cut at the patellar tendon to ensure proper tension on the Achilles tendon from the gastrocnemius muscle during fixation (n=3). Skin and toes were carefully removed to allow successful penetration of the fixative and decalcifying solutions. Hindlimbs were immersion-fixed in 10% neutral buffered formalin for 3 days and decalcified using Webb-Jee 14% EDTA for 14 days at room temperature. After routine paraffin processing, the hindlimbs were cut immediately below the knee, and the hind paw was carefully removed to avoid removing the calcaneus. Dissected hindlimbs were then paraffin-embedded. Sagittal sections (3 uM) were cut and baked overnight at 60°C. For immunofluorescence staining, sections were deparaffinized and rehydrated. Antigen retrieval (10mM sodium citrate, 0.05% Tween-20, pH 6.0) was performed at 75°C for 3 hours. Slides were blocked in blocking buffer (10% normal donkey serum [NDS] Jackson ImmunoResearch #017-000-121 in PBS + 0.1% Tween-20). Sections were then incubated with antibodies to alpha-smooth muscle actin (αSMA-FITC, 1:200, Sigma Aldrich #F3777) in blocking buffer overnight at 4°C. The following day, slides were washed with PBS for 3 x 5 minutes. Nuclei were counterstained with Hoechst 33342 (NucBlue™ Live ReadyProbes™ Reagent, Thermo Fisher Scientific), and slides were cover slipped (ProLong™ Diamond Antifade Mountant, Thermo Fisher Scientific). Imaging was performed using a VS120 Virtual Slide Microscope (Olympus).

### Cell Density Quantification

Fluorescent images of Hoechst 33342-stained VWR and sedentary control samples were processed using Visiopharm image analysis software v.2025.08.06 (Visiopharm). Regions of Interest (ROI) were drawn to include the entire Achilles tendon, as well as the insertion and midsubstance region, as described above (Transmission Electron Microscopy Imaging and Quantification). Hoechst 33342 positive staining was used to identify nuclei to quantify the total number of cells within each respective ROI. Automatic segmentation using a threshold classifier was used to define cells based on fluorescent intensity. Cell counts were normalized to the ROI’s total area in µm.

### Bulk-RNA Sequencing and Analysis

Achilles and plantaris tendons were isolated together from the hindlimb and snap-frozen in liquid nitrogen and stored at -80°C until analysis (n=7). Tendons from individual mice were homogenized in TRIzol reagent (Invitrogen #15596026) using a PowerMasher II (Nippi Inc. #891300). Total RNA was isolated using the RNeasy Plus Micro Kit (Qiagen #74034) per the manufacturer’s recommendations. The total RNA concentration was determined with the NanoDrop 1000 spectrophotometer, and RNA quality was assessed with the Agilent Bioanalyzer 2100. The average RNA integrity number (RIN) for all samples was 5.70 ± 0.58. Libraries were prepared and sequenced by the University of Rochester Genomics Research Center (UR GRC). 1 ng of total RNA was pre-amplified with the SMARTer Ultra Low Input kit v4 (Takara Bio #634895) per manufacturer’s recommendations. The quantity and quality of the subsequent cDNA were determined using the Qubit Fluorometer and the Agilent Bioanalyzer 2100. 150 pg of cDNA was used to generate Illumina-compatible sequencing libraries with the NexteraXT library preparation kit (Illumina #FC-131-1024) per the manufacturer’s recommendations. Amplified cDNA libraries were hybridized to the Illumina flow cell, and single-end reads were generated for each sample using Illumina NovaSeq6000. Raw reads generated from the Illumina base calls were demultiplexed using bcl2fastq version 2.19.1.

Differential expression analysis was performed using DESeq2-1.34.0 with an adjusted p-value threshold of 0.05. Heatmaps were created in R using Seurat v5 DoHeatmap() function. Bulk-RNA sequencing data will be made publicly available at GEO at the time of publication.

### Sample Preparation for Visium HD Spatial Transcriptomics

VWR and sedentary control mice were perfused-fixed with 4% paraformaldehyde (PFA). Hindlimbs were then dissected as described above. Hindlimbs were immersion-fixed in 4% PFA for 24 hours and decalcified in Morse’s solution (10% sodium citrate, 20% formic acid) for 48 hours at 4°C (n=5). Samples were submitted for routine paraffin processing. Sagittal (5 uM) sections were cut onto Nexterion Slide H – 3D Hydrogel Coated slides (Fisher Scientific, #NC0782819), baked at 42°C for 3 hours, and stored in a desiccator until further processing by the UR GRC. Sample preparation from this step onward was completed by the UR GRC. Samples were stained with Hematoxylin and Eosin (H&E) and imaged with a VS120 Virtual Slide Microscope (Olympus) to visualize overall tissue architecture. After imaging, samples were deparaffinized, destained, and decrosslinked. Tissue slides and Visium HD slides (10X Genomics, Pleasanton, CA) were loaded into the Visium CytAssist instrument, to be brought into proximity with one another. The Visium HD slide was removed from the Visium CytAssist for downstream library preparation. Spatial barcodes were used to associate reads with the tissue section images for spatial mapping of gene expression.

### Data Processing and Quality Control for Visium HD Spatial Transcriptomics

Space Ranger v4.0 (10X Genomics, Pleasanton, CA) was used for initial data processing and cell segmentation. The entire Achilles tendon (except for the myotendinous junction region where tissue damage occurred from sectioning) was manually annotated in Loupe Browser v9.0 (10X Genomics). Initial unsupervised clustering at resolution 0.3 had three distinct clusters in the VWR and control Achilles tendon (**Supplemental Fig. 1A, B**). A total of 1,751 and 1,383 cells were present in the VWR and control Achilles tendons, respectively (**Supplemental Fig. 1A**). Cells with low counts (fewer than 5) localized to the lateral portion in both VWR and control tendons, which aligned with cluster 2 cells (**Supplemental Fig. 1C-F**). Cluster 2 cells were then removed from analysis due to low counts, as low-count cells can significantly skew differential expression analysis and reduce gene detectability (41). After removing cluster 2, cells within the bottom 10% of total counts were removed during quality control, corresponding to cells with fewer than 4 counts in VWR samples and fewer than 5 counts in control samples. After filtering out the bottom 10% of cells based on low counts, a total of 1,553 and 1,210 cells remained in the VWR and control Achilles tendon, respectively (**Supplemental Fig. 1G, H**).

### Data Merging and Analysis for Visium HD Spatial Transcriptomics

All analysis for Visium HD spatial transcriptomic data was performed in Python using Squidpy and Scanpy (42). VWR and sedentary control samples were merged using the *anndata.concat* function. The merged data were normalized (*scanpy.pp.normalize_total*) and log-transformed (*scanpy.pp.log1p*). Dimensionality reduction was performed using principal component analysis (*scanpy.tl.pca*) followed by construction of a k-nearest neighbor graph (*scanpy.pp.neighbors*). A two-dimensional embedding generated using UMAP for visualization (*scanpy.tl.umap*). Cluster marker genes were identified with differential expression by comparing cells in each cluster to all remaining cells using the Wilcoxon rank-sum test (*scanpy.tl.rank_genes_groups*) function.

Scanpy.tl.rank_genes_groups was also used to identify differentially expressed genes (DEGs) between VWR and controls. DEGs had an adjusted p-value ≤ 0.5 with a fold change of greater than 0 in a natural log scale. Spatial transcriptomics data will be made publicly available at GEO at the time of publication.

### Statistical Analysis

Normality across all datasets for biomechanics, tendon cellularity, and normalized counts from bulk-RNA sequencing data was assessed using the Shapiro-Wilks test. Normal data were assessed using an unpaired Welch’s t-test (GraphPad Prism v.10.3). Body weight data were analyzed by multiple unpaired t-tests. Cellularity and nuclear aspect ratio data were analyzed by two-way ANOVA with multiple comparisons. Data that were not normal were assessed by a Mann-Whitney U Test (GraphPad Prism v.10.3). All data are shown as mean ± standard deviation, unless otherwise noted.

## Results

### Mice maintained consistent running levels and weight during eight weeks of VWR

To investigate tendon cell behavior at homeostasis, we first needed to establish a model of moderate exercise to simulate baseline tendon health. In skeletal and cardiac muscle, four weeks of VWR has previously shown to be sufficient to induce adaptations consistent with moderate exercise (35). Because tendons have a slow tissue turnover rate (43), we chose eight weeks of VWR to ensure sufficient time for the tendons to respond to load. To do so, we subjected mice to either VWR or normal cage activity (sedentary control; locked wheels) for eight weeks. Mice progressively increased their running distance in the first two weeks and plateaued at 5 weeks, running an average of 120.514 ± 19.60 km/week during weeks 5 through 8 (**Fig. 1A**). VWR mice ran a total distance of 902.88 ± 80.06 km during the eight week experiment (**Fig. 1B**). Control mice experienced significant weight gain (start weight: 26.86 ± 1.288g, final weight: 31.63 ± 2.912g, p=0.0055); however, VWR mice maintained their starting weight throughout (start weight: 26.19 ± 1.285g, final weight: 26.59 ± 0.902g, p>0.9999) (**Supplemental Fig. 2**).

**Figure 1.**
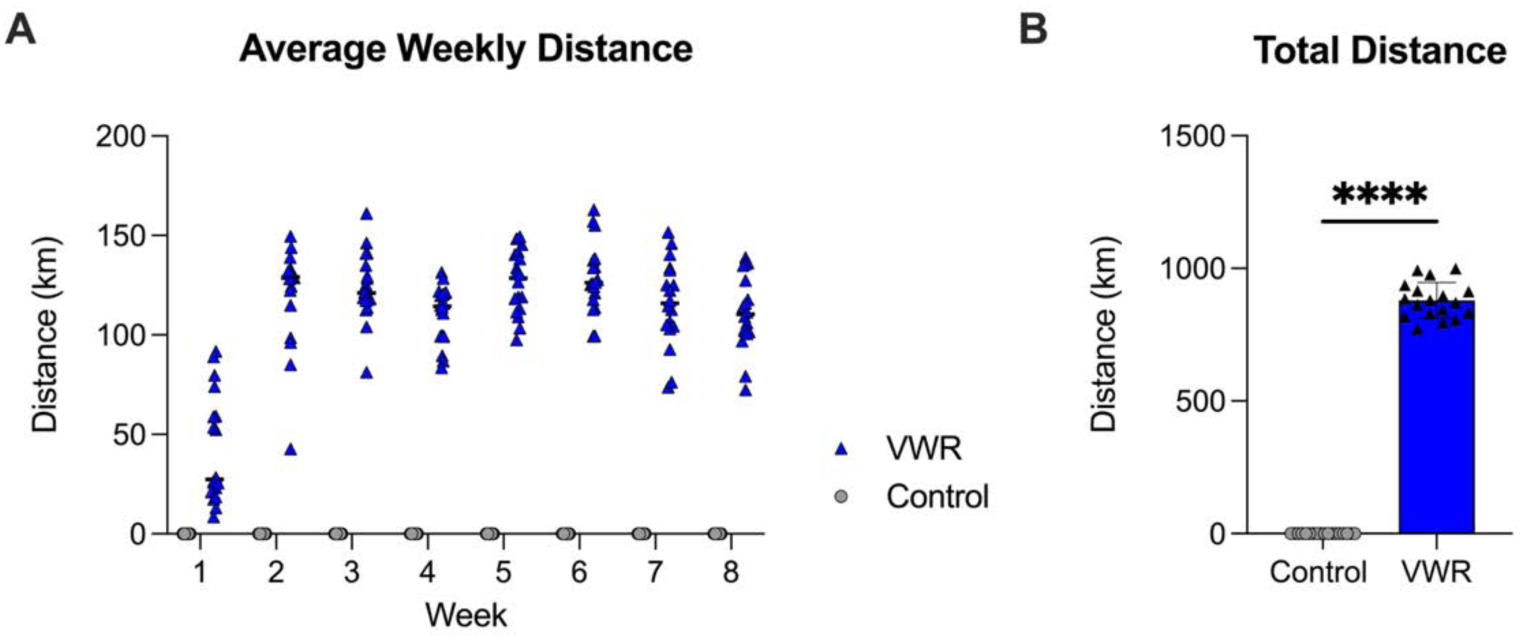
Average weekly and total running distance of VWR and control mice. (A) Average weekly running distance of control (gray) and VWR (dark blue) mice. (B) Average total running distance after 8 weeks between control (gray) and VWR (dark blue) mice. n = 19 mice per group. ****p≤0.0001.

### VWR improved the mechanical properties of Achilles tendons compared to controls

To assess functional changes after eight weeks of VWR, we performed uniaxial tensile testing at the Achilles tendon midsubstance. There was a significant decrease in the CSA between VWR and control tendons (control: 1.204 ± 0.302 mm^2^, VWR: 0.842 ± 0.120 mm^2^, p=0.0044) (**Fig. 2A**), indicating that VWR results in smaller tendons. There were no significant changes in the stiffness (control: 8.651± 1.342 N/mm, VWR: 8.984 ± 0.539 N/mm, p=0.4843) of VWR tendons compared to controls (**Fig. 2B**). Peak load (control: 14.940 ± 2.730 N, VWR: 17.470 ± 0.973 N, p=0.0178), peak stress (control: 12.800 ± 2.748 MPa, VWR:19.780 ± 4.700 MPa, p=0.0019), and elastic modulus (control: 31.070 ± 1.722 MPa, VWR: 44.680 ± 12.420 MPa, p=0.0117) were all significantly increased with VWR (**Fig. 2C, D, E**). We observed no significant changes in stress relaxation ratio (S: R) at 8% strain (control: 0.2561 ± 0.05920, VWR: 0.2928 ± 0.0404, p = 0.2029) or 16% strain (control: 0.2416 ± 0.0435, VWR: 0.2484 ± 0.3000, p = 0.8263) between VWR and controls (**Supplemental Fig. 3A, B**). No significant changes in the time to 50% relaxation at 8% strain (control: 6.020 ± 1.550 seconds, VWR: 5.967 ± 0.6205 seconds, p = 0.9218) or 16% strain (control: 6.970 ± 2.290 seconds, VWR: 6.678 ± 1.138 seconds, p=0.7263) were observed between VWR and controls (**Supplemental Fig. 3C, D**). Combined, these data indicate that the midsubstance is smaller and stronger, but not stiffer, in VWR Achilles tendons compared to controls.

**Figure 2.**
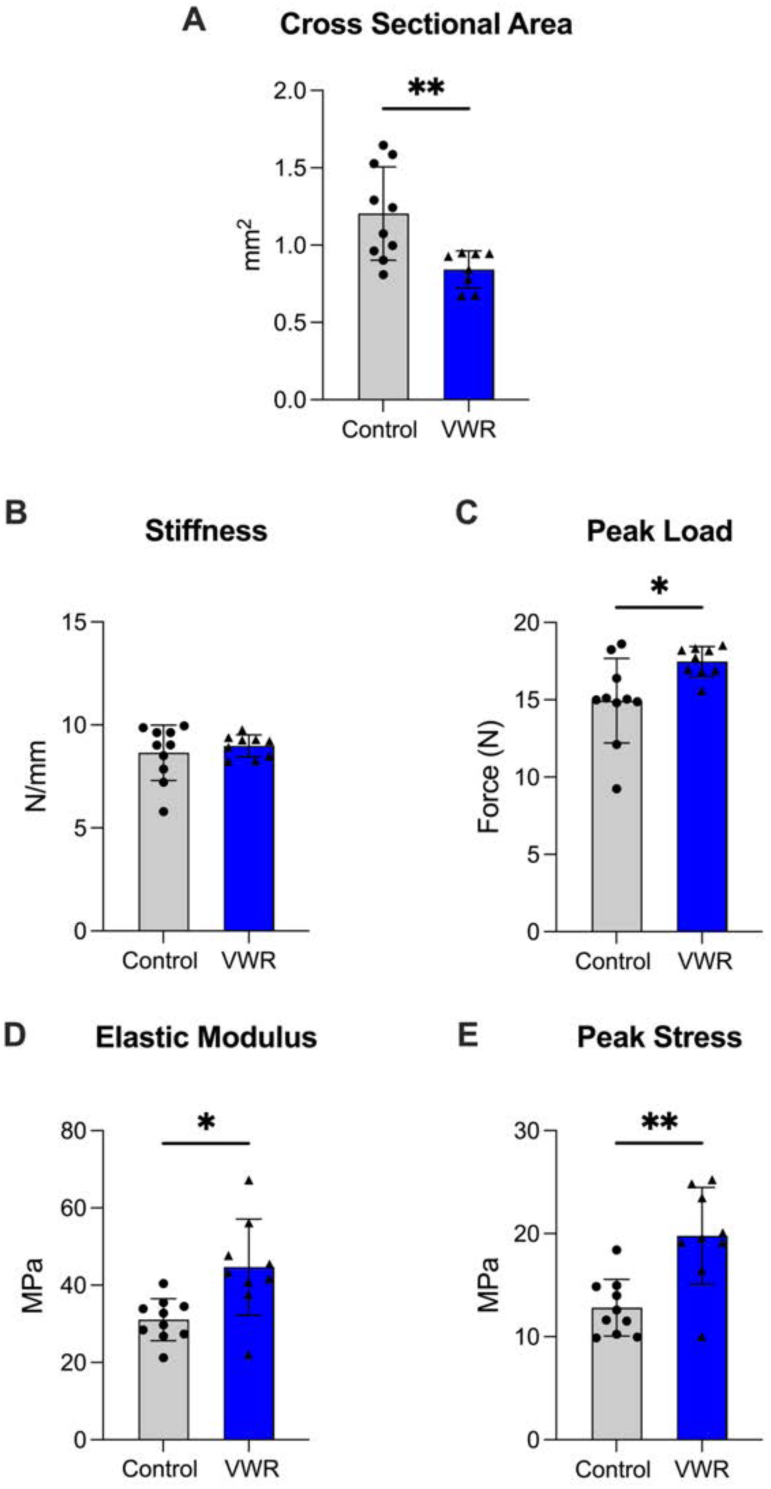
Biomechanical assessment at the midsubstance of the control and VWR Achilles tendons. Quantification of (A) Cross-Sectional Area, (B) Stiffness, (C) Peak Load, (D) Elastic Modulus, and (E) Peak Stress of control (gray) and VWR (blue) Achilles tendons. n=9-10 tendons from individual mice per group. *p≤0.05, **p≤0.01.

### VWR did not result in tendon degeneration or inflammation

As sustained mechanical loading can lead to tendon degeneration, we evaluated for the presence of cellular changes or alterations to the ECM that could indicate the presence of inflammation or damage with VWR. Given that we observed significant increases in the mechanical properties at the midsubstance, we divided subsequent analyses between the midsubstance and insertion to determine potential regional effects of VWR on tendon health. No differences in overall cellularity were observed in midsubstance (control: 4 × 10^−4^ ± 2.646 × 10^−5^ cells/µM^2^, VWR: 4 × 10^−4^ ± 5.292 × 10^−5^ cells/µM^2^, p=0.9999) or insertion (control: 6.267 × 10^−4^ ± 3.055 × 10^−5^ cells/µM^2^, VWR: 5.833 × 10^−4^ ± 1.801 × 10^−4^ cells/µM^2^, p=0.7191) (**Supplemental Fig. 4A, B**), suggesting no recruitment of inflammatory cells or proliferation of resident tendon cells. In both control and VWR tendons, cellularity was significantly decreased in the midsubstance compared to the insertion, indicating that the insertion maintains a higher cell density than the midsubstance, independent of VWR (control insertion: 6.267 × 10^−4^ ± 3.055 × 10^−5^ cells/µM^2^, control midsubstance: 4 × 10^−4^ ± 2.646 × 10^−5^ cells/µM^2^ p=0.0201; VWR insertion: 5.833 × 10^−4^ ± 1.801 × 10^−4^ cells/µM^2^, VWR midsubstance: 4 × 10^−4^ ± 5.292 × 10^−5^ cells/µM^2^, p=0.0475) (**Supplemental Fig. 4B**). There was no difference in cell nuclear aspect ratio in the midsubstance (control: 3.070 ± 0.219, VWR: 2.904 ± 0.214, p=0.3986) or the insertion (control: 2.741 ± 0.130, VWR: 2.634 ± 0.4219, p=0.7000) (**Supplemental Fig. 4A, B**). As tenocyte rounding is a hallmark of tendon degeneration (25, 44), this suggests that tenocyte health is not negatively impacted by eight weeks of VWR. We also observed no α-SMA+ cells that would indicate tenocyte activation to a myofibroblast phenotype in either the insertion or midsubstance of VWR tendons (**Supplemental Fig. 4C**). To monitor for damage to the collagen ECM with VWR, we stained with Collagen Hybridizing Peptide (CHP)-Cy3, which specifically identifies denatured collagen. No CHP staining was observed in either the insertion or midsubstance of VWR or control Achilles tendons (**Supplemental Fig. 4D**). Collectively, these data indicate that eight weeks of VWR does not result in damage or degeneration and suggest that the increased mechanical properties we observed at the midsubstance reflect healthy, adaptive remodeling rather than a degenerative response to mechanical loading.

### Expression of genes associated with collagen remodeling was increased with VWR

To investigate the transcriptional changes underlying the increased mechanical and material properties, we performed bulk-RNA sequencing on control and VWR Achilles tendons. VWR was associated with a modest transcriptional response, with 46 differentially expressed genes (DEGs; 22 down- and 24 up-regulated) compared to controls (**Fig. 3A**). Notably, several upregulated genes have established roles in collagen remodeling, such as lumican [*Lum*] (p=0.0028), fibroblast activation protein [*Fap*] (p=0.0008), and prolyl 4-hydroxylase subunit alpha 3 [*P4ha3*] (p=0.0003) (**Fig. 3B**). Consistent with prior reports of mechanical loading inducing changes in collagen organization (45, 46), these findings suggest that VWR results in increased collagen remodeling compared to controls.

**Figure 3.**
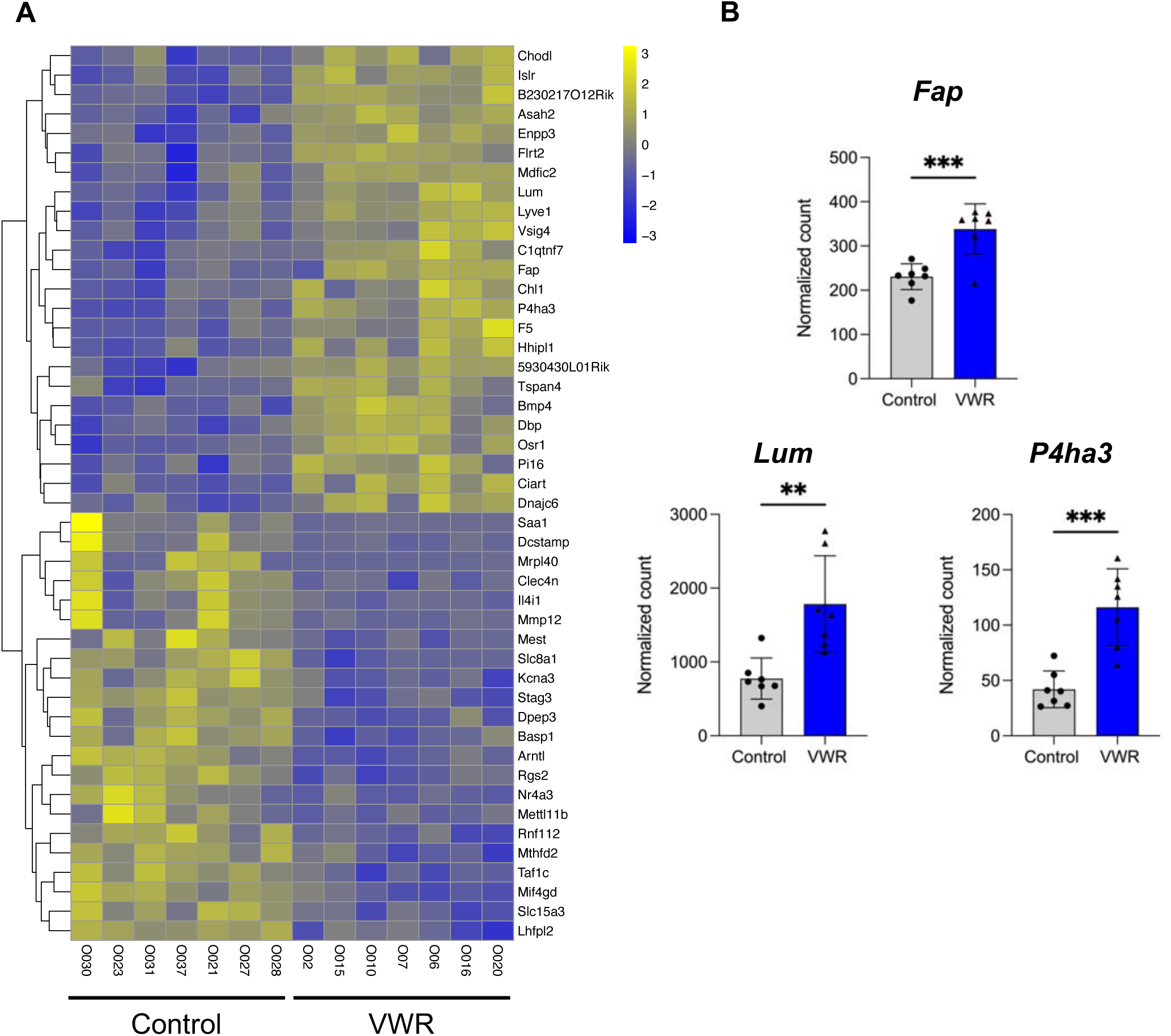
Bulk-RNA sequencing of VWR and control Achilles tendons. (A) Heatmap of all Differentially Expressed Genes (DEGs) between control and VWR Achilles tendons. (B) Normalized counts of Fibroblast Activation Protein (*Fap*), Lumican (*Lum*), and Prolyl 4-Hydroxylase Subunit Alpha 3 (*P4ha3*) between control (gray) and VWR (blue) Achilles tendons from bulk-RNA sequencing data. n=7 tendons from individual mice per group. **p≤0.01,***p≤0.001.

### VWR resulted in a shift towards larger collagen fibrils in the midsubstance

Next, to determine if the decreased CSA and increased biomechanical properties at the midsubstance with VWR corresponded to structural changes in the collagen matrix, we performed Transmission Electron Microscopy (TEM) in a region-specific manner (insertion vs. midsubstance). No significant differences were observed in the collagen fibril density in either the insertion (control: 58.687 ± 6.3182, VWR: 58.088 ± 4.4282, p=0.8669) or midsubstance (control: 57.615 ± 8.047, VWR: 56.334 ± 5.264, p=0.7746) with VWR (**Fig. 4A, B**). Binned analysis of fibril diameters (Diameter 1: 0-60 nm, Diameter 2: 60-120nm, Diameter 3: 120-180 nm, Diameter 4: >180 nm) also revealed no significant differences in the insertion between control and VWR (**Fig. 4C, D**). At the midsubstance, the percent of D1 fibrils was significantly decreased with VWR compared to controls (control = 16.144 ± 3.504%, VWR= 10.806 ± 2.116%, p=0.0484) (**Fig. 4E, F**). Although not statistically significant, the percent of D3 fibrils was increased with VWR compared to controls (control: 27.649 ± 2.903%, VWR: 33.068 ± 5.119%, p=0.0891) (**Fig. 4F**). No significant differences were observed in the percent of D2 fibrils (control = 22.714 ± 2.167%, VWR = 21.967 ± 1.840%, p=0.6185), or D4 fibrils (control: 33.492 ± 4.271%, VWR = 32.781 ± 6.613%, p=0.8634) (**Fig. 4F**). Taken together, these findings suggest that while there are no overall differences in collagen fibril size at the insertion, there is a significant decrease in small (D1) collagen fibrils and a non-significant shift towards larger (D3) collagen fibrils at the midsubstance following VWR.

**Figure 4.**
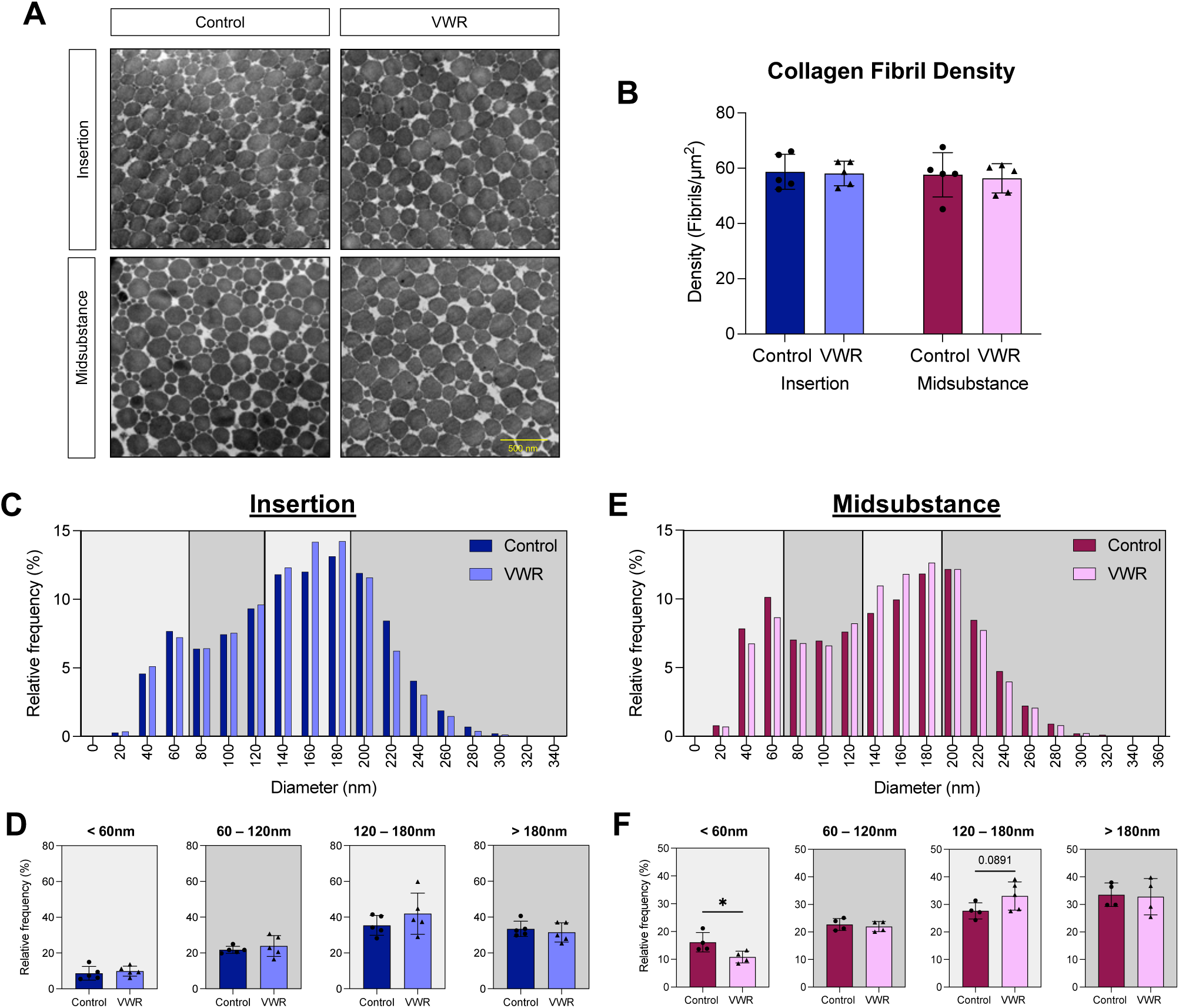
Transmission Electron Microscopy (TEM) data of collagen fibrils in VWR and control Achilles tendons. (A) Representative TEM images of collagen fibrils in the insertion (top) and midsubstance (bottom) between control (left) and VWR (right) tendons. Scale bar = 500 nm. (B) Quantification of collagen fibril density in the insertion (blue) and midsubstance (pink) between VWR and controls. Histogram of collagen fibril frequency distribution and binned analysis of different diameters (D1:<60 nm, D2: 60-120 nm, D3: 120-180 nm, D4: >180 nm) at the (C, D) insertion and (E, F) midsubstance between VWR and controls. n=4-5 tendons from individual mice per group. *p<0.05.

### Achilles tendons exhibited region-specific baseline transcriptional profiles

Since we detected significant changes following VWR at the midsubstance (biomechanics and TEM), but not at the insertion, we leveraged Visium HD spatial transcriptomics (47) to determine the transcriptional response to physiological load at a single-cell level in a region-specific manner. Unsupervised clustering of all cells in the VWR and control Achilles tendon identified two distinct clusters (0 and 1) (**Fig. 5A**). Spatial mapping of these clusters onto H&E-stained sections of VWR and control Achilles tendons revealed that cluster 0 localized to the midsubstance while cluster 1 localized to the calcaneal insertion (**Fig. 5B**). Cluster 0 was the predominant population in both VWR and control tendons (72.53% vs. 61.89%), whereas cluster 1 comprised a smaller fraction of cells (27.47% vs. 38.11%) (**Fig. 5C**). Differential expression analysis between these two clusters revealed distinct transcriptomic signatures (**Fig. 5D**). Top genes in both clusters 0 and 1 included genes related to secreted matrix proteins [*Dcn* (48), *Clec3a* (49)] and the glycoprotein *Clu* (50), suggesting that these genes are expressed broadly throughout the tendon, rather than exhibiting region-specific expression (**Fig. 5D**).

**Figure 5.**
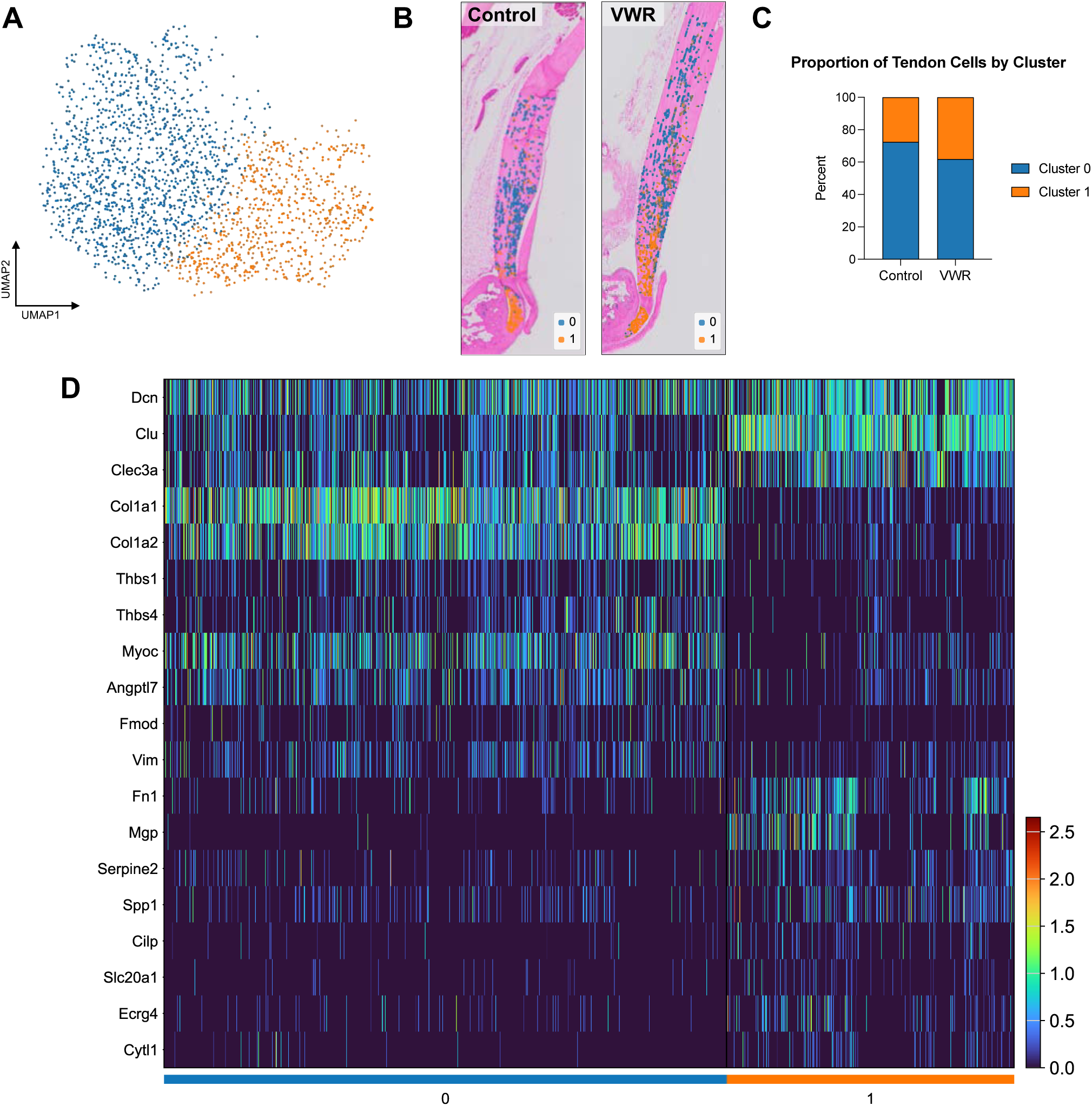
Spatial differential expression analysis of VWR and control Achilles tendon cells. (A) Uniform Manifold Approximation and Projection (UMAP) of unsupervised clustering at resolution 0.3 of merged filtered cells from VWR and control Achilles tendons. (B) Clusters 0 (blue) and 1 (orange) overlaid onto the control (left) and VWR (right) Achilles tendon H&E image. (C) Percent distribution of clusters 0 (blue) and 1 (orange) between VWR and controls. (D) Heatmap of top DEGs from clusters 0 (blue) and 1 (orange).

Compared to cluster 1 (insertion) cells, cluster 0 (midsubstance) had upregulated expression of type 1 collagen genes [*Col1a1*, *Col1a2* (3)], matricellular thrombospondin family genes [*Thbs1, Thbs4* (51)], trabecular network-associated genes [*Myoc* (52), *Angptl7* (53)], small leucine-rich proteoglycan [*Fmod* (54)] and intermediate filament gene [*Vim* (55)] (**Fig. 5D**). In contrast, cluster 1 (insertion) was enriched for matrix remodeling genes [*Fn1* (56), *Serpine2* (57)], cartilage homeostasis [*Cilp* (58), *Cytl1* (59), *Ecrg* (60)] and mineralization-associated [*Mgp* (61)*, Spp1* (62)*, Scl20a1* (63)] genes compared to cluster 0 cells (**Fig. 5D**). Taken together, these data suggest that cluster 0 (midsubstance) and cluster 1 (insertion) cells have distinct transcriptional profiles that relate to the function of the specific regions. Specifically, cluster 0 cells express genes consistent with the fibrillar collagen-rich midsubstance. Interestingly, genes associated with the trabecular meshwork of the cornea [*Myoc, Angptl7*] were also identified, suggesting that these genes may represent previously unrecognized components of the midsubstance. In contrast, cluster 1 cells express genes related to maintaining the fibrocartilage interface at the insertion.

### Achilles tendons displayed region-specific transcriptional responses to VWR

To determine if cluster 0 (midsubstance) and cluster 1 (insertion) have distinct responses to physiological load, we performed differential expression analysis comparing VWR cluster 0 and 1 to their corresponding control clusters. In response to VWR, cluster 0 (midsubstance) cells upregulated matrix remodeling [*Comp*, *Thbs4]*, trabecular meshwork network [*Myoc*], and secreted protein [*Clu*, *Ccdc80*] related genes compared to cluster 1 cells (**Fig. 6A**). At the midsubstance, upregulation of *Comp* and *Thbs4* at the midsubstance is consistent with enhanced matrix remodeling in response to physiological load (51, 64) (**Fig. 6B, E**). *Clu*, a glycoprotein and secreted chaperone involved in protection against oxidative stress (50), may contribute to mediating exercise-induced stress responses at the midsubstance (**Fig. 6C**). To our knowledge, *Clu*, *Myoc,* and *Ccdc80* have no reported roles in tendon (**Fig. 6C, D, F**). In contrast, cluster 1 (insertion) cells upregulated bone-[*Spp1* (*62*)], and cartilage-associated [*Chad* (65)] genes with VWR, suggesting maintenance of the fibrocartilage interface (**Fig. 7A, B, C**). The load-induced matricellular gene *Sparc* was also upregulated in cluster 1 (insertion) cells compared to controls, consistent with previous studies demonstrating that *Sparc* may play a key role in the tendon mechanoresponse (7) (**Fig. 7D**). Notably, *Comp*, which is known to support collagen fibrillogenesis (66, 67), was upregulated in both clusters with VWR, suggesting a role in maintaining tendon homeostasis independent of region (**Fig. 6A** and **Fig. 7A**). Combined, these data indicate that cells in cluster 0 (midsubstance) and cluster 1 (insertion) upregulate distinct sets of matrix remodeling genes that reinforce the collagen-rich midsubstance and fibrocartilaginous insertion identity, respectively.

**Figure 6.**
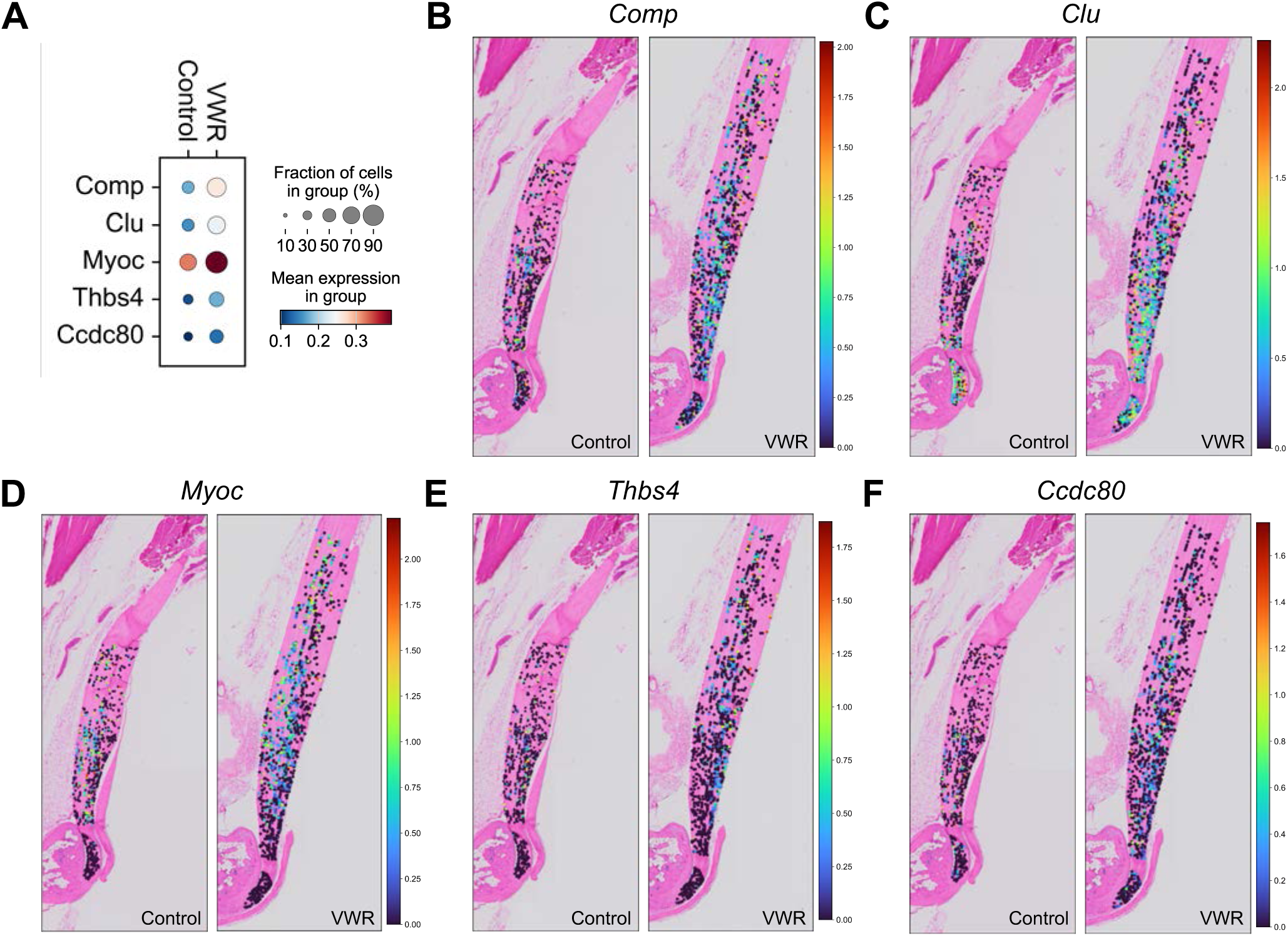
Top genes upregulated with VWR in the midsubstance compared to control (cluster 0). (A) Dot plot of the top upregulated genes in cluster 0 cells between VWR and control cells. Spatial mapping of (B) *Comp*, (C) *Clu*, (D) *Myoc,* (E) *Thbs4*, and (F) *Ccdc80* expression on H&E image of VWR and control Achilles tendons.

**Figure 7.**
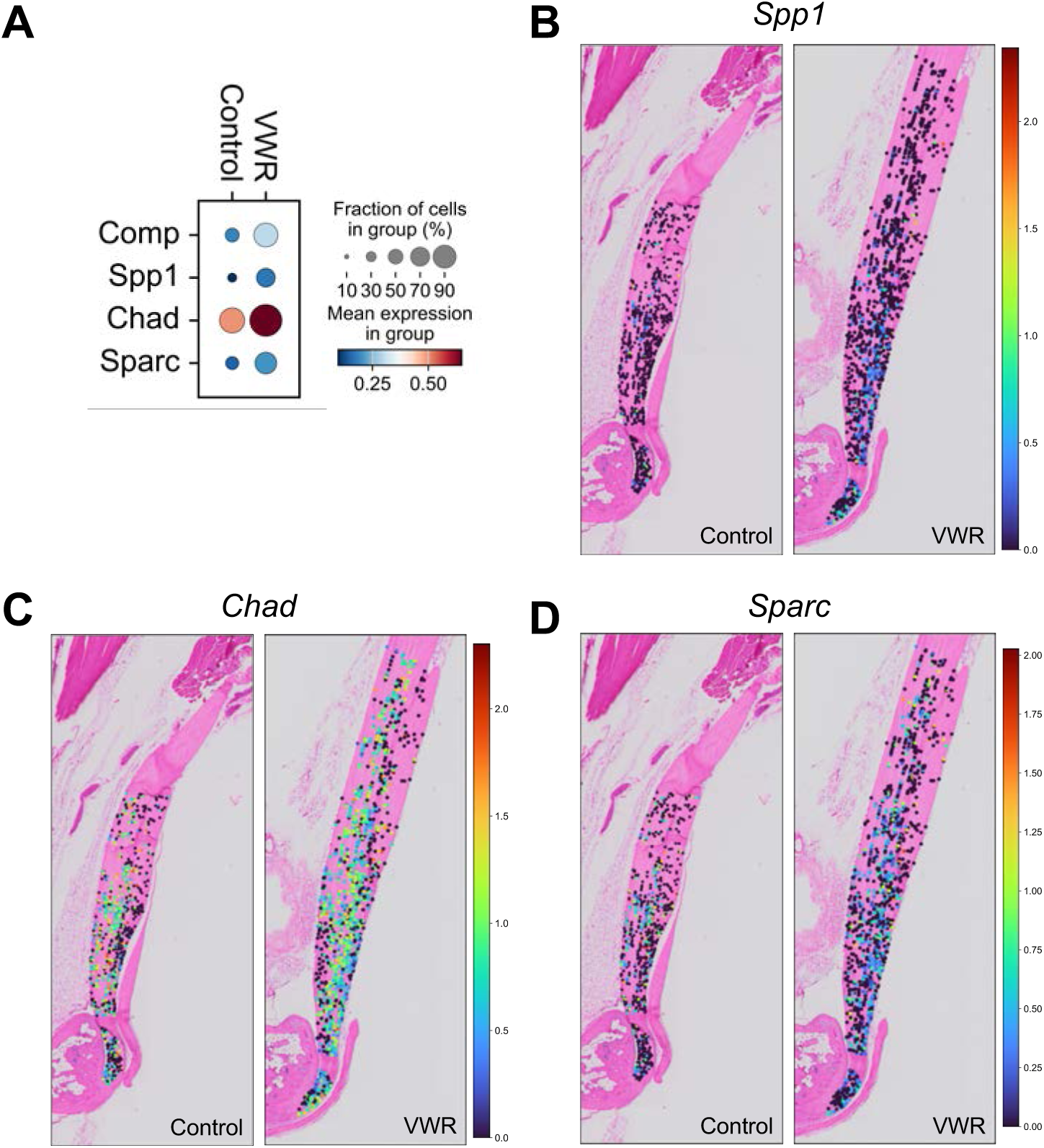
Top genes upregulated with VWR in the insertion compared to control (cluster 1). (A) Dot plot of the top-upregulated genes in cluster 1 cells between VWR and control cells. Spatial mapping of (B) *Comp*, (C) *Spp1*, (D) *Chad,* (E) *Sparc* expression on H&E image of VWR and control Achilles tendons.

## Discussion

Failure to maintain tendon homeostasis compromises tissue integrity and mechanical properties, leading to impaired mobility of affected individuals. Understanding the fundamental cellular contributions to tendon homeostasis in response to physiological load is therefore critical to identifying potential targets to maintain tendon health. Here, we report an in-depth characterization of the tendon cellular response using a VWR model. Eight weeks of VWR resulted in smaller and stronger, but not stiffer, Achilles tendons at the midsubstance, compared to controls. While increased tendon stiffness can be considered a beneficial adaptation during healing that can enhance force transmission (68, 69), excessive stiffening may impair function (70). Importantly, we found no evidence of tendon degeneration indicative of overuse in Achilles tendons following eight weeks of VWR. We observed no collagen damage with VWR, suggesting preservation of tendon collagen structure. Despite increased upregulation of *Fap* in VWR Achilles tendons compared to controls by bulk RNA-seq, we did not detect myofibroblasts, which are typically associated with fibrotic matrix remodeling. This suggests that fibroblast activation in this context may reflect alternative roles, such as extracellular matrix production and organization, rather than a pro-fibrotic response (71, 72). Further, no changes in tendon cellularity were observed, which is in contrast to the increased cell density typically associated with an influx of immune cells during fibrotic healing of tendons (73). By spatial RNA-seq, *Spp1* (osteopontin) was upregulated with VWR at the insertion, compared to controls. *Spp1* is well characterized in bone, where it coordinates osteoblast and osteoclast activity to maintain homeostasis, suggesting that increased *Spp1* expression at the insertion may similarly support maintenance and remodeling of the fibrocartilage structure in response to physiological load. Consistent with this role in ECM regulation, *Spp1* knockout mice have significantly increased collagen fibril diameter and reduced density in patellar tendons compared to wild-type controls (74). Moreover, following loss of mechanical load via denervation, *Spp1* knockout mice have larger collagen fibrils with increased density in patellar tendons, compared to wild-type controls (74). Although *Spp1* is frequently associated with inflammation (75) and fibrosis (76, 77), our findings suggest that, in the context of physiological loading, increased *Spp1* expression is more likely to reflect ECM remodeling and turnover rather than a pathological response. Taken together, our data indicated that VWR represents a physiological loading regimen, rather than pathological loading or overuse.

Increased expression of genes associated with collagen remodeling and a shift towards larger collagen fibrils in the tendon midsubstance suggest that collagen remodeling is a primary driver underlying the tendon response to VWR, but that this occurs in a region-specific manner. Correspondingly, analysis by spatial transcriptomics indicated that cells in the insertion and midsubstance have distinct transcriptional responses to VWR. Specifically, our data suggest maintenance of fibrocartilage and collagen fibril-rich interface of the insertion and midsubstance, respectively. We also identified uncharacterized genes in tendon [*Myoc*, *Clu*, *Ccdc80*] at the midsubstance, where increased biomechanical properties were observed. Investigating how *Myoc*, *Clu*, and *Ccdc80* contribute to homeostasis will provide new insights into the cellular processes underlying the tendon cell response to physiological load. Collectively, these data show region-specific cellular responses to physiological loading in the mouse Achilles tendon that result in tissue-level adaptive changes. Given the prevalence of region-specific tendinopathies (e.g., insertional (78, 79) and midsubstance (80, 81) tendinopathies), understanding region-specific cellular responses to physiological load can help inspire potential therapeutic targets for promoting tendon health.

Surprisingly, eight weeks of VWR led to a significant reduction in Achilles tendon midsubstance CSA. Despite this decrease, tendons exhibited superior biomechanical properties at the midsubstance following VWR. To our knowledge, this is the first study evaluating VWR on male C57BL/6J mouse Achilles tendon homeostasis. A recent study using female high-runner mice selectively bred for increased voluntary running activity (∼3-fold greater running distance compared to controls) reported enhanced biomechanical properties following eight weeks of wheel running compared to non-high-runner and sedentary controls, despite no differences in CSA between groups (82). While both studies found stronger tendons with VWR, differences in strain, age, and sex limit direct comparison between studies. Other loading models like FTR report no differences or increased CSA (8), but these results likely reflect overuse rather than physiological loading. Collectively, these studies suggest that different loading regimens, duration, and anatomical regions likely have distinct effects on CSA. Importantly, whether these changes in CSA represent stable structural changes or transient responses to load remains unclear.

TEM analysis results did not reveal smaller collagen fibrils that might be expected with a reduced CSA. In fact, we observed a significant decrease in small collagen fibrils and a shift towards larger collagen fibrils, which may underlie the increased mechanical and material properties at the midsubstance. These structural changes are further supported by our spatial transcriptomics data, in which midsubstance cells upregulate non-collagenous, matricellular genes such as *Comp* (66), *Thbs4* (51), and *Dcn* (83), all of which are known regulators of collagen fibrillogenesis and ECM organization. Thus, changes in the non-collagenous matrix could also contribute to the reduced midsubstance CSA through increased matrix compaction or reorganization. Additionally, we did not assess water content with VWR in this study, which could impact tendon CSA and biomechanical properties following physiological loading (84, 85).

Spatial transcriptomics data further support maintenance of homeostasis in response to physiological load, with region-specific cellular responses that preserve the specialized structure of the Achilles tendon. Specifically, cells in the midsubstance upregulated genes associated with fibrillar collagen remodeling, whereas cells in the insertion upregulated fibrocartilage-associated genes, consistent with maintenance of their distinct structural and functional identities in response to physiological load. Notably, we also identified genes with previously uncharacterized roles in tendon, including clusterin (*Clu)*, myocilin (*Myoc),* and coiled-coil domain containing 80 (*Ccdc80)* that are upregulated following VWR. Interestingly, a prior proteomic analysis of blood plasma from VWR mice (four weeks) identified clusterin as upregulated compared to sedentary controls (86). Recombinant clusterin decreased inflammatory gene expression and attenuated memory loss in a mouse model of Alzheimer’s, suggesting a potential exercise-induced protective role (86). Clusterin and myocillin are also highly expressed in the trabecular meshwork of ocular tissues where they regulate intraocular pressure (87, 88). Although their roles in tendon homeostasis are unknown, previous work has shown that myocilin can bind to extracellular matrix proteins such as collagens, fibronectin, and decorin, and regulate metalloprotease activity (89). Similarly, clusterin has been implicated in regulating the trabecular meshwork extracellular matrix and actin cytoskeleton (88, 90). Taken together, these findings suggest that clusterin and myocilin may contribute to extracellular matrix remodeling at the midsubstance. Future work will focus on identifying functional roles for clusterin and myocilin in maintaining tendon homeostasis in response to physiological load.

*Ccdc80* was also upregulated specifically in the midsubstance with VWR. *Ccdc80* is a secreted protein that has been reported to be expressed in mouse Achilles tendons (91). Interestingly, *Ccdc80* was identified as an exerkine that is produced by human skeletal muscle in response to exercise and protects against cardiac fibrosis and hypertrophy (92). *Ccdc80* knockout mice have increased sensitivity to high-fat diet-induced glucose intolerance and reduced glucose-stimulated insulin secretion, suggesting that Ccdc80 may also regulate glucose and energy homeostasis (93). Given its enrichment in the midsubstance region and its known roles in metabolic regulation and exercise, these findings suggest that Ccdc80 may play a metabolic role in maintaining tendon homeostasis at the midsubstance in response to physiological load. Future work will define the role that Ccdc80 plays in tendon homeostasis through knockout studies.

Laboratory mice, such as C57BL/6J mice used in this study, have decreased activity as compared to wild mice (94). Wild mice are known to move extensively within their home range, with distances of up to 20 km a day (95, 96). This suggests that normal cage activity in laboratory mice limits their natural activity levels, resulting in relatively low habitual tendon loading. VWR may therefore provide a stimulus that maintains tendon health by mimicking the mouse’s natural behavior. Collectively, our findings support a model in which Achilles tendons in standard laboratory conditions may exist in an underloaded state, and that physiological load via VWR induces positive extracellular matrix remodeling that enhances mechanical function without inducing tendon damage.

### Limitations

The initial goal of this study was to define the precise cellular processes that underpin tendon homeostasis in response to physiological load. However, a major limitation of current spatial transcriptomic technologies is the low sequencing depth. In contrast to bulk or single-cell RNA sequencing, where RNA is efficiently extracted from lysed cells, spatial approaches capture RNA *in situ* on a thin (5 µM) formalin-fixed paraffin-embedded (FFPE) section. Sample preparation for FFPE sections, including fixation, decalcification, and paraffin processing, can further degrade RNA, reducing transcript recovery. In addition, Visium HD spatial transcriptomics employs a probe-based capture system, which constrains detection to transcripts that efficiently hybridize to probes. As a result, highly expressed and stable transcripts, such as ECM-related genes, are more readily detected, whereas low-abundant, transiently expressed transcripts like transcription factors are more likely to drop out. This technical bias towards ECM-related genes likely contributes to the limited overlap between our spatial and bulk RNA sequencing datasets. Although we were unable to identify specific transcription factors or cellular processes that maintain homeostasis in response to physiological load, we were able to identify region-specific transcriptional responses at a single-cell level. Future work will focus on integrating single-cell sequencing data with the spatial transcriptomics data to investigate cellular contributions to homeostasis in response to physiological load with higher resolution while maintaining spatial context.

Another limitation of this study is the exclusive use of male mice. Because VWR requires single housing to accurately measure running distance, males were selected due to their more stable behavioral and physiological responses under these conditions (38). Female mice also exhibit greater variability in running activity, due in part to their estrus cycle (35). Prior studies have reported that male C57BL/6J Achilles tendons are larger, with reduced cellularity and lower expression of ECM-related proteins such as periostin, fibronectin, and tenascin-c, compared to females (97). Similarly, in humans, male Achilles tendons have larger CSA and increased stiffness than those of females (98). Together, these findings suggest that Achilles tendons have sex-dependent differences in structure and function and may respond differently to physiological load to maintain homeostasis, which will be of future investigation.

This study also evaluated only one time point and therefore cannot resolve whether earlier changes potentially precede the structural and functional changes seen at eight weeks, or if these changes persist following cessation of running. It is also unclear whether prolonged loading (beyond eight weeks) is also beneficial, or if there is a point at which it eventually becomes maladaptive. Future studies will aim to clarify both the preceding cellular events that result in beneficial changes at eight weeks and the long-term effects of VWR on Achilles tendon health.

Finally, we only tested biomechanical properties at the midsubstance and not at the insertion. Because the insertion is structurally and compositionally distinct from the midsubstance, the region-specific cellular responses observed with spatial transcriptomics do not directly translate to the observed increases in midsubstance mechanical and material properties. Future studies incorporating region-specific mechanical testing at the insertion will be important for linking these transcriptional changes to functional tissue properties.

### Conclusions

In summary, eight weeks of physiological loading via VWR resulted in smaller but mechanically superior tendons at the midsubstance, with a corresponding decrease in the proportion of small fibrils and a shift towards an increased proportion of large fibrils. While increased frequency of large fibrils supports the functional phenotype observed at the midsubstance, our spatial transcriptomic data suggest that changes in the non-collagenous matrix could be driving changes in CSA. With spatial transcriptomics, we determined that the insertion and midsubstance have region-specific cellular responses to physiological load, with upregulated genes promoting maintenance of the fibrocartilage and collagen-rich matrix, respectively. Although our analysis was limited by the sequencing depth inherent to spatial transcriptomics, we identified genes with previously unknown roles in tendon homeostasis (*Myoc*, *Clu* and *Ccdc80*), underscoring the power of discovery-based transcriptomics approaches to understand fundamental tendon biology. Ultimately, understanding the precise cell-specific mechanisms underlying homeostasis in a region-specific manner will be important for identifying ways to promote tendon health.

## Acknowledgements

The authors thank the Center for Musculoskeletal Research Histology, Biochemistry and Molecular Imaging (HBMI) core for technical assistance with preparing samples for histology and the UR Genomics Research Core for assistance with bulk- and spatial- RNA sequencing. We would also like to extend our sincere thanks to the Shriner’s Children Hospital’s Multi-Imaging Core (Portland, OR) for their technical assistance with the transmission electron microscopy imaging.

## Author Contributions

Conceived and designed research: SM and AECN, performed experiments: SM, EL, NS, AK, AECN, analyzed data: SM, EL, EK, AK, interpreted results of experiments: SM, MB, AECN, prepared figures: SM and AECN, drafted manuscript: SM, edited and revised manuscript: SM and AECN, approved final version of manuscript: SM, EL, NS, AK, EK, MB, AECN.

## Supplemental Figures

**Supplemental Figure 1.**
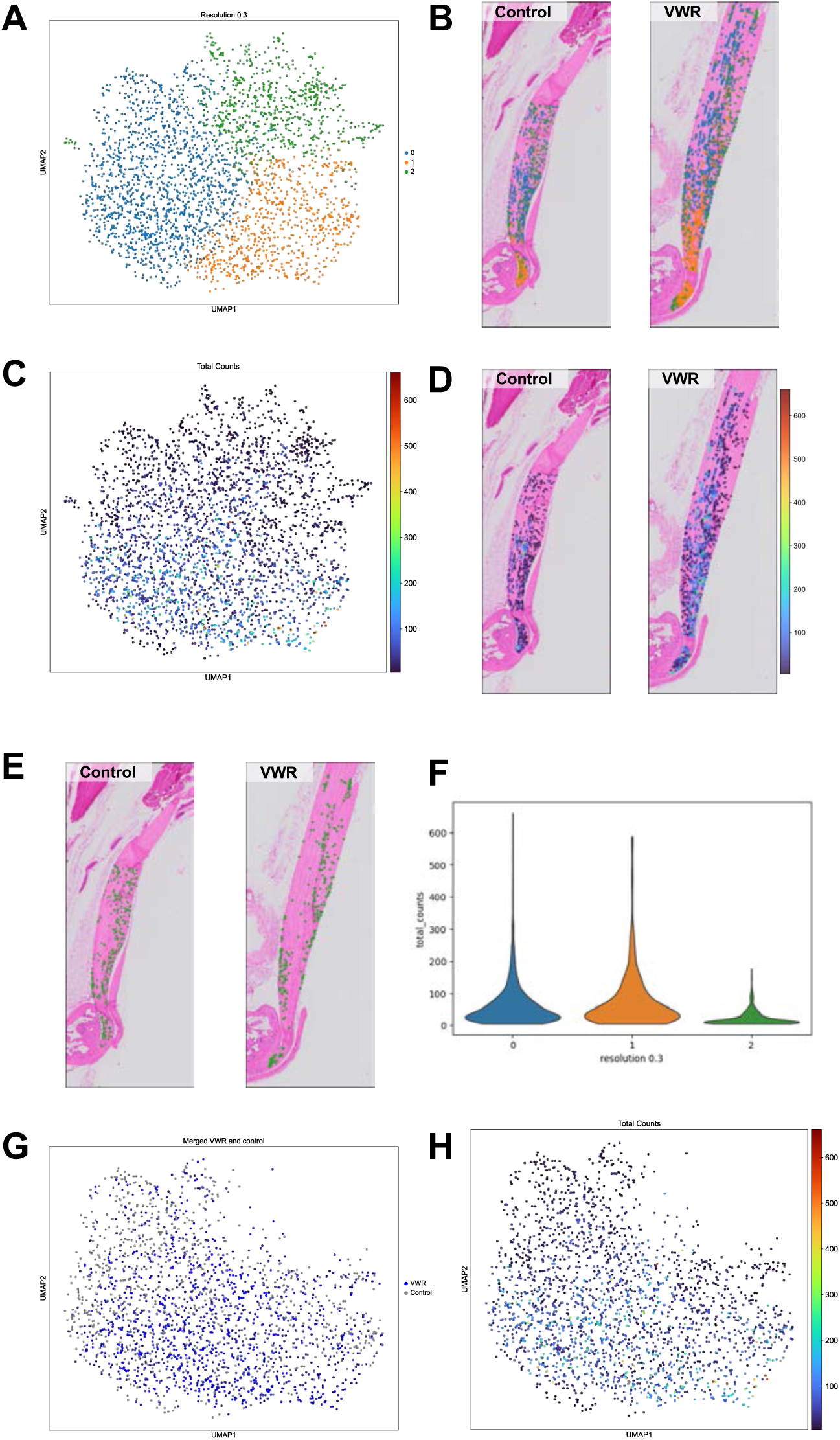
Quality control assessment of tendon cells in spatial transcriptomics data. (A) Uniform Manifold Approximation and Projection (UMAP) of unsupervised clustering at resolution 0.3 of merged VWR and control data. (B) Spatial mapping of clusters 0-2 onto the Hematoxylin and Eosin (H&E) image of VWR and control Achilles tendons. (C) UMAP of total counts in VWR and control Achilles tendon cells. (D) Spatial mapping of total count data on H&E images of VWR and control Achilles tendons. (E) Cluster 2 cells on H&E image of VWR and control Achilles tendons. (F) Violin plot of total counts in each cluster of VWR and control Achilles tendons. (G) UMAP of merged VWR (blue) and control (gray) Achilles tendon cells after filtering out cells within the bottom 10% of total counts. (H) UMAP of total counts after filtering out cells within the bottom 10% of total counts. n=1 per group.

**Supplemental Figure 2.**
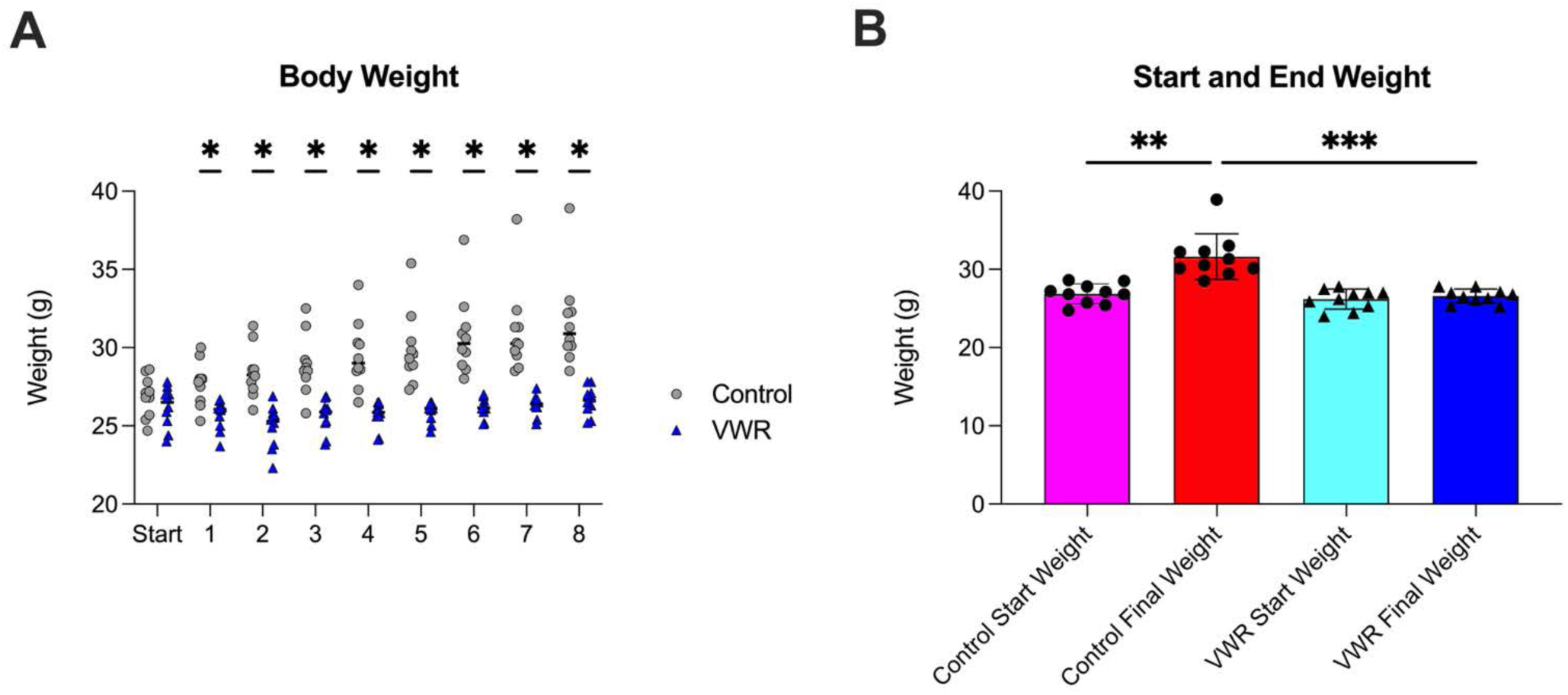
Mouse body weight data of VWR and control Achilles tendons. (A) Weekly body weight (g) of VWR (blue) and control (gray) mice. (B) Start and end weight for control (pink and red bars) and VWR mice (light and dark blue bars). n=9-10 mice per group. *p≤0.05, **p≤0.01, ***p≤0.001.

**Supplemental Figure 3.**
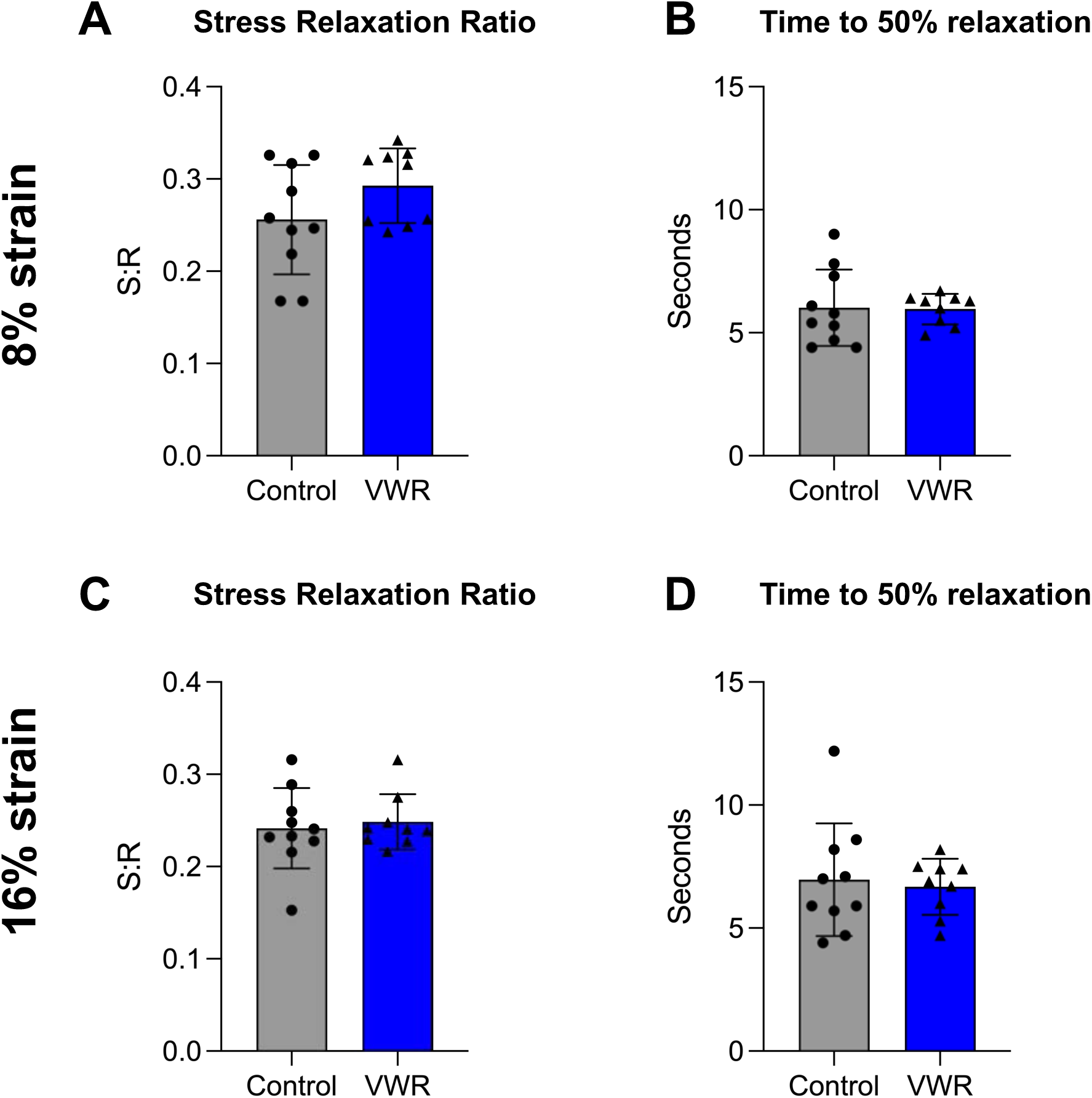
Stress relaxation data of VWR and control Achilles tendons. (A, C) Stress Relaxation ratio (S:R) and (B,D) time to 50% relaxation at 8% strain (top) and 16% strain (bottom). n=9-10 mice per group.

**Supplemental Figure 4.**
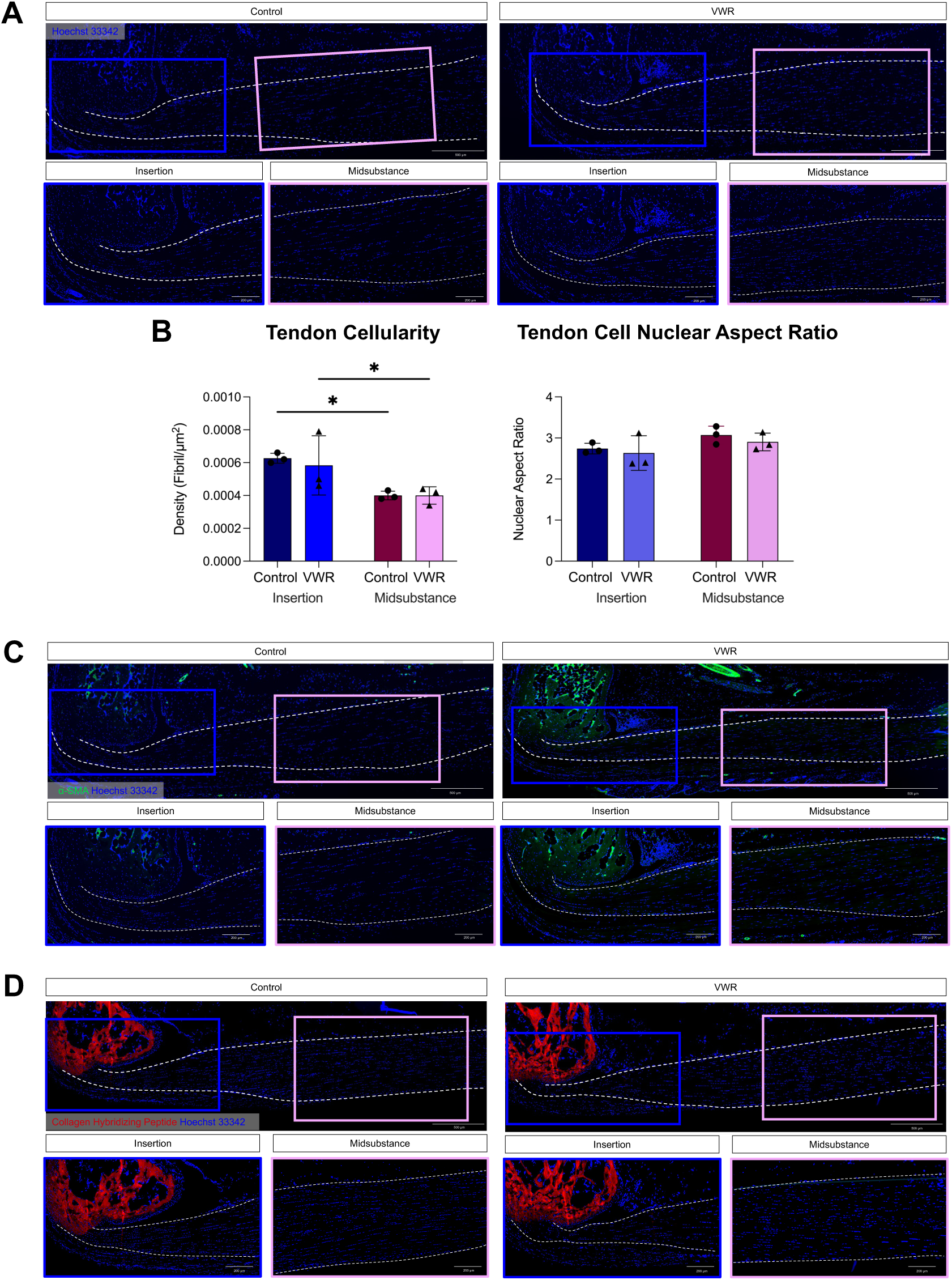
Hoechst 33342, α-SMA, and Collagen Hybridizing Peptide (CHP) staining of VWR and control Achilles tendons. (A) (Top) Representative images of control and VWR Achilles tendons stained with Hoechst 33341. (Bottom) Magnified view of insertion (blue) and midsubstance (pink). (B) Quantification of total cell density (left) and nuclear aspect ratio (right) in the insertion and midsubstance. (C) (Top) Representative images of control and VWR Achilles tendons stained with α-SMA FITC and Hoechst 33341. (Bottom) Magnified view of insertion and midsubstance. (D) (Top) Representative images of control and VWR Achilles tendons stained with CHP-Cy3 and Hoechst 33341. (Bottom) Magnified view of insertion and midsubstance. Achilles tendons are outlined by white dashed lines. Scale bars = 500 µm, magnified image scale bars = 200 µm. n=3-4 per group. *p≤0.05.

